# Sister species, shared and divergent responses: early transcriptomic profiling of *Bursaphelenchus xylophilus* exposed to *Esteya* spp

**DOI:** 10.64898/2026.10.11.758299

**Authors:** David Pires, Ana Rita Varela, Miroslav Kolařík, Maria L. Inácio, Manuel Mota, Cláudia S. L. Vicente

## Abstract

*Bursaphelenchus xylophilus* (pinewood nematode, PWN) responds transcriptionally to nematophagous fungi, but the earliest events preceding cuticle penetration are poorly characterized, and whether closely related fungal antagonists elicit comparable responses is unknown. Here, we compared PWN transcriptomic responses to *Esteya floridanum* V14639 (EF) and *E. vermicola* CCF3117 (EV) at 1 h post-inoculation (hpi), capturing the onset of host–fungus interaction. Despite their phylogenetic proximity, under our exposure conditions the two fungi elicited partly divergent host responses at 1 hpi. EF induced a broad transcriptional response enriched in peptidase activity and lysosomal pathways, accompanied by downregulation of cuticle structural components. In contrast, EV triggered a narrower response, without the proteolytic and lysosomal enrichment observed with EF. Direct comparison of the two treatments revealed no significantly enriched Gene Ontology categories, although pathway analysis identified DNA replication and repair pathways, comprising genes more highly expressed under EF. Enrichment analyses revealed a shared transcriptional signature in both interactions characterized by downregulation of xenobiotic- and central carbon-metabolism pathways, suggesting an early reduction in xenobiotic detoxification capacity rather than induction of detoxification programs. These findings suggest that phylogenetic proximity does not necessarily predict conserved early transcriptional responses, although comparisons of single isolates cannot separate species-level from isolate-level effects.

**Graphical abstract:** 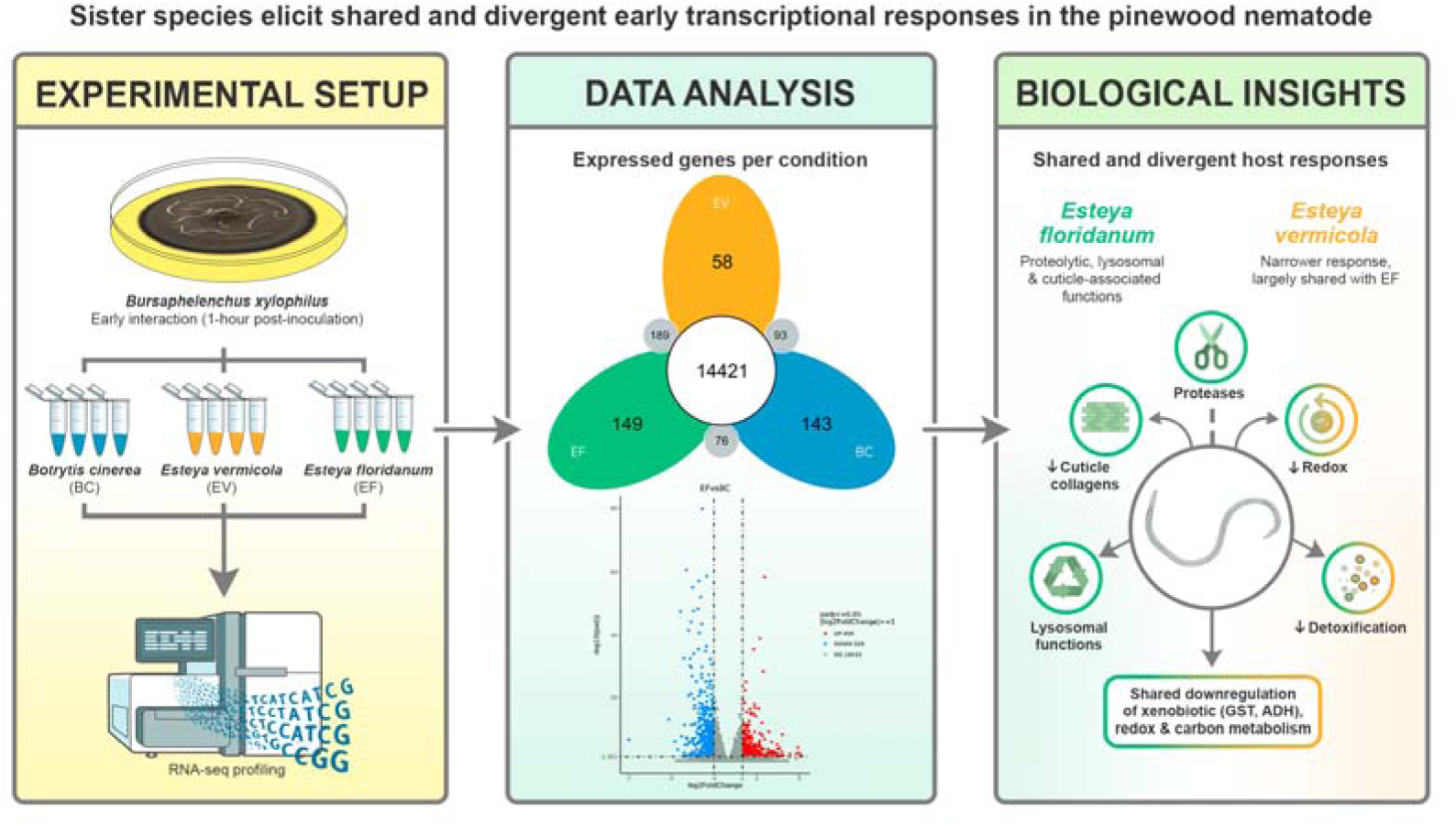

## Introduction

Closely related pathogens do not necessarily elicit conserved host responses; yet, this assumption underlies many biocontrol deployment strategies, including those targeting the pinewood nematode (PWN), *Bursaphelenchus xylophilus*, a quarantine pest vectored by *Monochamus* spp. and responsible for pine wilt disease (PWD) [1]. Coniferous forests face a significant and expanding biosecurity threat from PWD, with traditional management approaches yielding limited success and insecticide-based vector control carrying non-target ecological costs [2–4]. In this context, nature-based pest management within a One Health framework has gained prominence through microbial antagonists that offer sustainable alternatives [5–7]. This approach is particularly urgent in Southern Europe, where PWD has caused widespread forest loss [8].

The success of fungal biocontrol agents depends critically on their ability to overcome host defenses, yet the molecular determinants of host susceptibility remain poorly predictable. Understanding early host-pathogen interactions could inform strain selection and deployment strategies, particularly for closely related antagonist species that may trigger markedly different defensive responses despite similar taxonomic classification. Nematode responses to fungal pathogens involve conserved innate immune pathways that have been extensively characterized in *Caenorhabditis elegans*, including pathogen recognition receptor signaling, antimicrobial peptide production, and cuticle remodeling [9–11]. However, these defense mechanisms can vary substantially across nematode species and ecological contexts. Recent work has shown that *C. elegans* responds to nematode-trapping fungi by downregulating cuticle collagens to reduce adhesion [12], while other nematodes may favor active structural reinforcement strategies. The extent to which *B. xylophilus* employs similar or divergent defense strategies against nematophagous fungi remains unknown, despite its economic importance as a forest pest.

*Esteya* spp. are well-studied fungal antagonists of the PWN and have shown considerable potential for biocontrol [13]. Both *E. vermicola* and *E. floridanum* have demonstrated nematophagous activity, and recent *in planta* experiments showed that preemptive inoculation with either species significantly reduced *B. xylophilus* reproduction in *P. pinaster* seedlings, whereas effects on symptom severity, disease progression, and survival were not consistently observed under the conditions tested [14]. These findings highlight the importance of understanding the mechanisms underlying *Esteya*–PWN interactions. *Esteya* spp. infect the PWN via lunate spores that attach to the cuticle and penetrate host tissues [15–20]. Molecular studies show that *E. vermicola* produces volatile organic compounds mimicking pine host signals to attract nematodes [21], harbors serine proteases for cuticle penetration [22], and associates with bacterial endosymbionts that may enhance virulence [23,24]. Unlike *E. vermicola*, which has been studied extensively at the molecular level, *E. floridanum* was only recently described and its interaction mechanisms with the PWN remain largely uncharacterized, making direct molecular comparison particularly informative [25]. Nevertheless, the early molecular events governing host responses and the extent to which different *Esteya* species elicit distinct early host transcriptional responses remain unknown.

To address this gap, we conducted a transcriptomic analysis of *B. xylophilus* exposed to *E. vermicola* (EV) and *E. floridanum* (EF), relative to nematodes maintained on their standard laboratory food fungus, *Botrytis cinerea* (BC), specifically targeting the 1-hour post-inoculation (hpi) window. Because *Esteya* pathogenesis follows an ordered chronological sequence (spore adhesion, germ tube penetration and subsequent colonization), molecular events occurring prior to cuticular breach are of particular interest. We hypothesize that EV and EF will elicit distinct transcriptional programs during the early interaction phase, reflecting differences in their early modes of host interaction. By restricting sampling to this early interaction phase, this study avoids the confounding transcriptional noise associated with the deteriorating physiological states and systemic tissue damage typical of later infection stages reported for EV [17]. Consequently, we provide comparative, high-resolution insights into early host transcriptional response dynamics in the *Esteya*–PWN interaction. More broadly, this work addresses whether early host responses to fungal sister species are conserved or can diverge into distinct interaction strategies.

## Results

### Differential gene expression and enrichment analyses

Differential expression analysis relative to the BC control (adjusted *P*-value, *P_adj_* ≤ 0.05, absolute log_2_ fold change, |log_2_FC| ≥ 1) revealed strongly divergent nematode responses in magnitude (Fig. 1A–B) and functional composition (Fig. 2). The PWN exhibited a strong transcriptional response to EF, with 459 upregulated and 529 downregulated differentially expressed genes (DEGs), whereas the response to EV was more limited, with 194 upregulated and 305 downregulated DEGs. Overall, EF elicited nearly twice as many DEGs as EV (988 vs 499), with the difference most pronounced among upregulated DEGs (2.4-fold; 459 vs 194). At the whole-transcriptome level, the three conditions shared a large common set of 14,421 expressed genes, with comparatively few genes detected exclusively in the EF (149), EV (58), or BC (143) samples (Fig. 1C).

**Figure 1.**
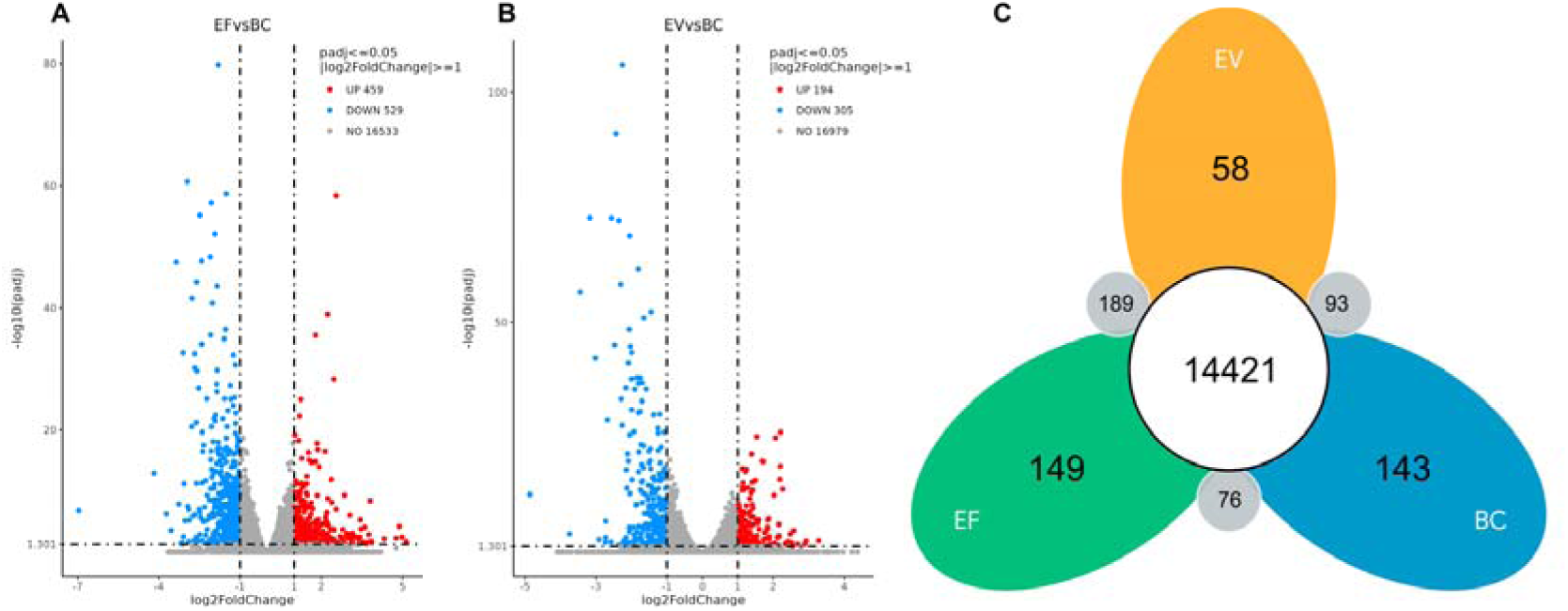
Differential gene expression in *Bursaphelenchus xylophilus* exposed to *Esteya* species. Volcano plots showing differentially expressed genes (DEGs) with *P_adj_* ≤ 0.05 and |log_2_FC| ≥ 1 for (A) *E. floridanum* vs *Botrytis cinerea* control (EF vs BC) and (B) *E. vermicola* vs *B. cinerea* control (EV vs BC). Red, blue and grey dots represent upregulated, downregulated, and non-significant genes, respectively. (C) Petal diagram showing the overlap of expressed genes among the three conditions. The central value indicates genes expressed in all conditions, petal values indicate genes expressed exclusively in one condition, and values in the small circles indicate genes shared between two conditions only.

**Figure 2.**
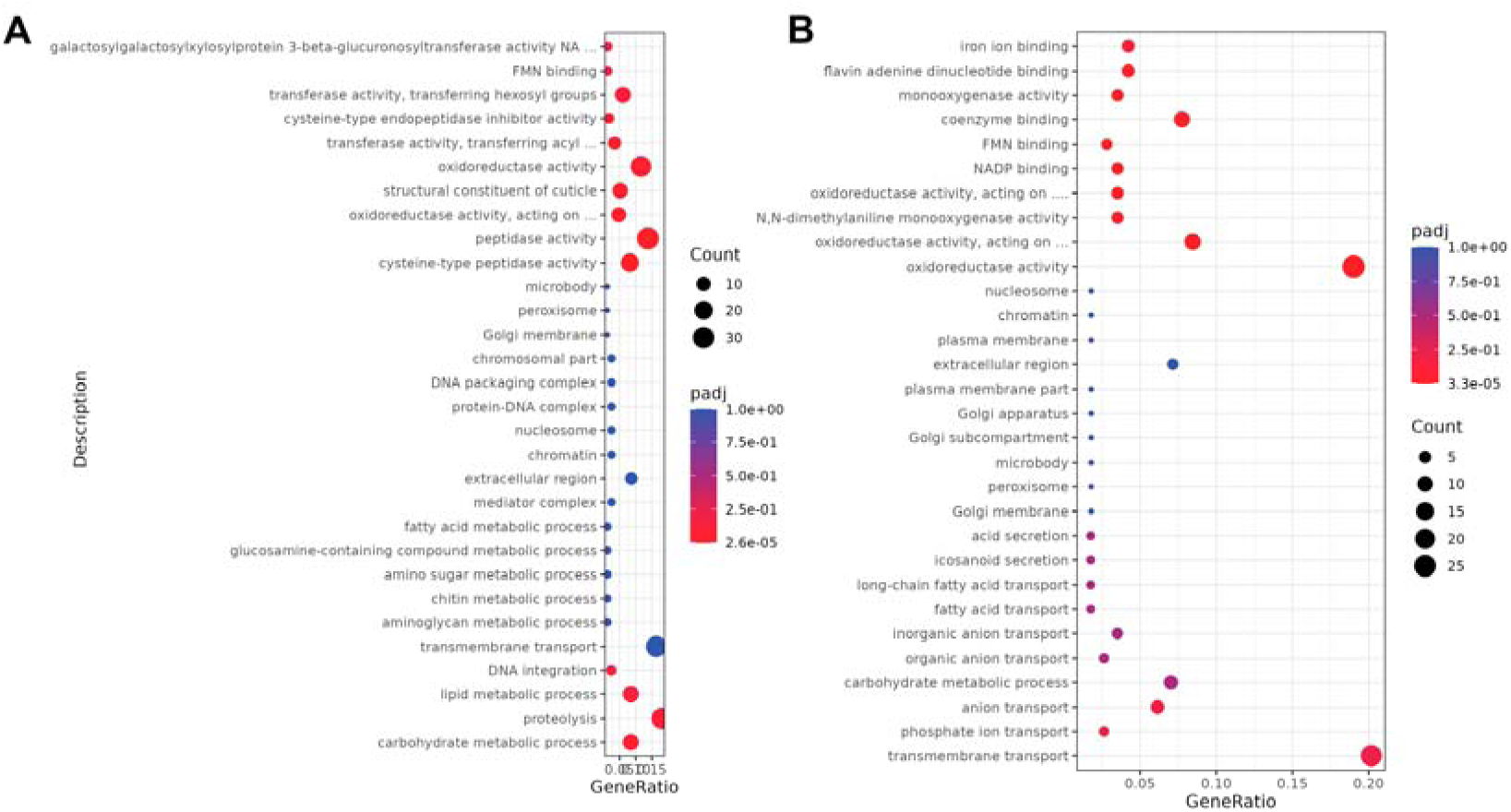
Gene Ontology (GO) enrichment in *Bursaphelenchus xylophilus* exposed to *Esteya* species. Top-ranked GO terms for (A) *E. floridanum* vs *Botrytis cinerea* control (EF vs BC) and (B) *E. vermicola* vs *B. cinerea* control (EV vs BC). Terms with *P_adj_* < 0.05 were considered significantly enriched. Dot size represents the number of DEGs annotated to each term, and color represents the adjusted *P*-value; color scales differ between panels.

Exposure to EF was associated with enrichment of proteolysis- and cuticle-associated functions (Fig. 2A). Gene Ontology (GO) enrichment analysis revealed highly significant enrichment of “cysteine-type peptidase activity” (GO:0008234, n = 19 genes, *P_adj_* = 2.57×10^−5^) and “peptidase activity” (GO:0008233, n = 32 genes, *P_adj_*= 0.017). Consistently, “cysteine-type endopeptidase inhibitor activity” (GO:0004869) was also enriched, comprising exclusively downregulated genes (4/4; *P_adj_* = 0.049). The EF interaction was further associated with enrichment of “structural constituent of cuticle” (GO:0042302, n = 12 genes, *P_adj_* = 0.033), comprising exclusively downregulated cuticle collagens (12/12 genes downregulated). In contrast, the EV interaction showed no significant GO enrichment for proteolytic or cuticle-related functions (Fig. 2B). The EV response was characterized by enrichment of metabolic and redox-associated functions, with the most significant GO enrichment being “oxidoreductase activity” (GO:0016491, n = 27 genes, *P_adj_* = 3.34×10^−5^), driven by monooxygenases, NADP-binding enzymes, and flavin-dependent oxidoreductases involved in redox homeostasis.

Because EF and EV were tested in separate exposures, their responses were compared by contrasting the independent enrichment results obtained for each against BC (Fig. 2A–B). Both interactions shared enrichment of “oxidoreductase activity” (GO:0016491) and “oxidoreductase activity, acting on paired donors, with incorporation or reduction of molecular oxygen” (GO:0016705), representing a shared oxidoreductase signature. Both terms were predominantly composed of downregulated genes and reached lower adjusted *P*-values in EV (GO:0016491: 25/27 genes downregulated, *P_adj_* = 3.34×10^−5^; GO:0016705: 11/12, *P_adj_* = 9.5×10^−5^) than in EF (GO:0016491: 21/27, *P_adj_* = 0.041; GO:0016705: 7/11, *P_adj_* = 0.033). Direct comparison of EV and EF yielded no significantly enriched GO terms (Supplementary Fig. S1).

Kyoto Encyclopedia of Genes and Genomes (KEGG) pathway analysis strongly corroborated the GO enrichment patterns and revealed additional mechanistic insights (Fig. 3). EF responses showed highly significant enrichment of lysosomal pathways (cel04142, n = 26 genes, 14 upregulated and 12 downregulated, *P_adj_* = 4.24×10^−6^) (Fig. 3A). Lysosome was not significantly enriched in EV (*P_adj_* = 0.12). EV responses were enriched for carbon metabolism (cel01200; 7/8 genes downregulated, *P_adj_* = 0.048) and fatty acid degradation (cel00071; 5/6 downregulated, *P_adj_*= 0.041), pathways that showed the same predominantly downregulated pattern in EF but narrowly missed significance (*P_adj_* = 0.058 and 0.051, respectively). Both interactions shared significant enrichment of glycolysis/gluconeogenesis (cel00010; EF: 11/12 genes downregulated, *P_adj_* = 1.49×10^−4^; EV: 8/9, *P_adj_*= 6.29×10^−4^) and pyruvate metabolism (cel00620; EF: 9/10, *P_adj_*= 2.05×10^−3^; EV: 7/8, *P_adj_* = 2.26×10^−3^) (Fig. 3A–B). Both interactions also showed enrichment of xenobiotic-metabolism pathways, including “drug metabolism – cytochrome P450” (cel00982) and “metabolism of xenobiotics by cytochrome P450” (cel00980), predominantly composed of downregulated genes (Fig. 3A–B). EF reached lower adjusted *P*-values for both pathways (cel00982: 20 genes, 16 downregulated, *P_adj_* = 2.23×10^−9^; cel00980: 19 genes, 15 downregulated, *P_adj_* = 2.88×10^−9^) than EV (cel00982: 14 genes, 12 downregulated, *P_adj_* = 5.91×10^−7^; cel00980: 11 genes, 8 downregulated, *P_adj_* = 8.63×10^−5^), although these were among the most significantly enriched pathways in both interactions. In both interactions, the DEGs mapped to these pathways were glutathione S-transferases (GST), alcohol dehydrogenases (ADH), flavin-containing monooxygenases, UDP-glucuronosyltransferases (UGTs) and an epoxide hydrolase, rather than cytochrome P450 genes; ten were shared between EF and EV, nine of which were downregulated in both. Outside these pathways, six cytochrome P450 genes were differentially expressed in both conditions, five of which were downregulated in both (BXY_1622200, BXY_1577300, BXY_0411400, BXY_0817800, and BXY_0170100). Direct comparison of the two treatments (EV vs EF) identified 141 DEGs, of which 118 (84%) were more highly expressed under EF. Although no GO categories were significantly enriched, KEGG analysis identified three significantly enriched pathways: nucleotide excision repair, base excision repair, and DNA replication (all *P_adj_* = 0.020), comprising exclusively genes expressed at higher levels under EF than EV (Supplementary Fig. S1).

**Figure 3.**
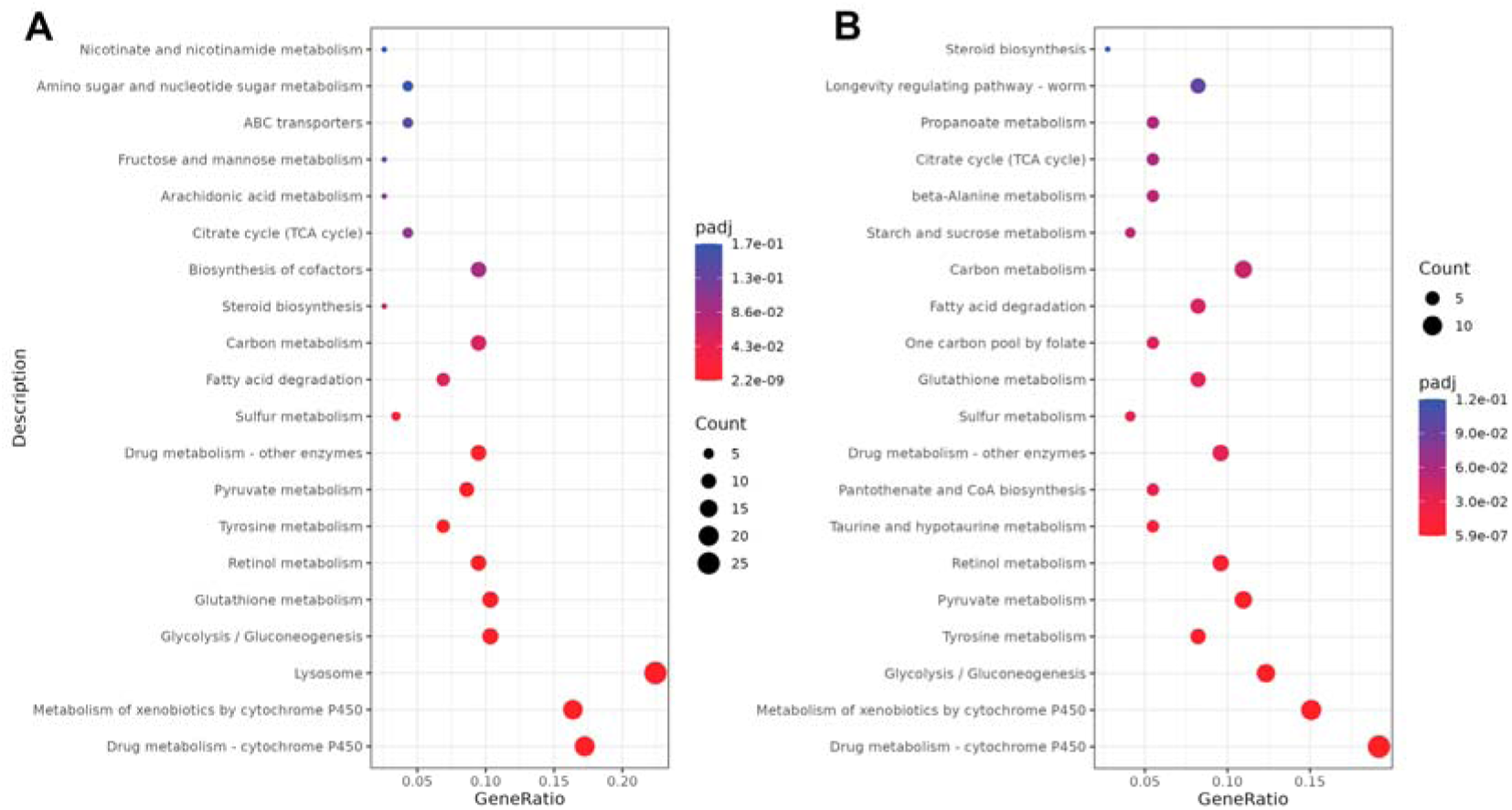
KEGG pathway enrichment in *Bursaphelenchus xylophilus* exposed to *Esteya* species. Top-ranked KEGG pathways for (A) *E. floridanum* vs *Botrytis cinerea* control (EF vs BC) and (B) *E. vermicola* vs *B. cinerea* control (EV vs BC). Pathways with *P_adj_* < 0.05 were considered significantly enriched. Dot size represents the number of DEGs annotated to each pathway, and color represents the adjusted *P*-value; color scales differ between panels.

### Validation of differential gene expression by RT-qPCR

Reverse transcription quantitative PCR (RT-qPCR) analysis of seven selected DEGs (BXY_0225800, BXY_1167200, BXY_1196500, BXY_0492900, BXY_0475400, BXY_1160600 and BXY_0936900) showed the same direction of change as RNA-seq for seven of the ten contrasts tested (Supplementary Table S2). These included the flavin-containing monooxygenase BXY_1167200 in both the EV vs BC and EV vs EF contrasts, and the upregulation of BXY_0936900 in EF.

## Discussion

Pathogen sensing of host arrival represents a fundamental biological challenge that shapes early interaction dynamics. Pathogens generally must detect host-derived signals to trigger energetically costly developmental transitions only after confirming host presence, requiring recognition systems that are both reliable and robust [26,27]. Recognizing damage-associated molecular patterns is central to this ecological interplay, with nematode-derived ascarosides serving as kairomones for fungal predators [28,29]. To clarify how these interactions unfold at the molecular level, we characterized the early transcriptional responses of *B. xylophilus* exposed to nematophagous fungi of the *Esteya* genus at the onset of infection.

By standardizing fungal colony size and inoculum density, and sampling at 1 hpi, we targeted the early interaction phase, sampling within a window in which spore adhesion and initial recognition are expected to dominate [17], although infection was not verified microscopically. Sampling at this interval is intended to reduce confounding effects of systemic tissue damage typical of later infection stages.

Our data reveal markedly different responses of *B. xylophilus* to EF and EV, suggesting that early host transcriptional responses are not necessarily conserved between fungal sister species [18,25]. Despite their divergent transcriptional profiles, both *Esteya* species elicited a shared early response characterized by predominant downregulation of genes associated with xenobiotic metabolism. EF additionally elicited a broader transcriptional program involving proteolysis, lysosomal functions, and coordinated cuticle-associated functions, which was absent in EV. Exposure to EF altered the expression of cysteine-type peptidases, most of which were upregulated, and lysosomal pathways were significantly enriched (KEGG: cel04142, uniquely enriched in EF), although with a mixed direction of expression among lysosome-associated genes. Similar transcriptional enrichment of lysosome-associated pathways and proteolytic systems has been reported when *B. xylophilus* is challenged by the nematophagous fungus *Arthrobotrys robusta* [30]. Lysosomal functions have likewise been identified as stress-responsive in *B. xylophilus* under chemical challenge, where disruption of lipid homeostasis is accompanied by lysosomal and autophagy involvement [31]. More recently, host-derived monoterpenes were shown to activate lysosomal pathways in *B. xylophilus*, with pharmacological inhibition and silencing demonstrating functional roles for lysosomal activity in stress adaptation and resource allocation [32]. Together, these findings support lysosomal recruitment as a broader stress-responsive feature of PWN physiology, although lysosomal enrichment alone does not distinguish degradative defense from metabolic adaptation. The transcriptional profile we report in the present study is therefore consistent with remodeling of intracellular degradative processes during fungal challenge, potentially targeting invading structures or secreted effectors. Concurrently, the PWN downregulated a coherent set of cuticle collagens, suggestive of cuticle remodeling that may reduce adhesion rather than reinforce the barrier. This parallels *C. elegans*, which evades trapping fungi by downregulating cuticle collagens to reduce fungal adhesion [12], suggesting that *B. xylophilus* may employ a similar anti-adhesion strategy at the cuticle surface. Together with transcriptional enrichment of proteolysis- and lysosome-associated genes, our results are consistent with a two-pronged early response: modulation of proteolytic and lysosomal functions alongside reduced cuticular adhesion rather than structural reinforcement. In *B. xylophilus* challenged by *A. robusta*, cuticle and collagen were induced only at later time points, coincident with epidermal degradation [30].

In contrast, the response to EV was narrower, lacking the proteolytic, lysosomal and cuticle-associated enrichment observed following EF exposure. Its dominant features, downregulation of oxidoreductase activity and of central carbon and fatty acid metabolism, were not specific to EV but formed part of a metabolic signature shared with EF, albeit with weaker statistical support for some pathways in EF. The EV response therefore appears largely nested within the EF response rather than constituting a distinct program. Whether this reflects attenuated host responsiveness or differences in interaction kinetics requires further experimental validation. Transcriptional induction of oxidative and detoxification pathways is a hallmark of *B. xylophilus* responses to chemical stressors, including fungal metabolites and bioactive compounds, where detoxification pathways dominate over structural defenses [33,34].

Consistent with the shared oxidoreductase signature identified by GO enrichment, both interactions also shared a common enriched core of KEGG pathways, including the xenobiotic-metabolism pathways “drug metabolism – cytochrome P450” and “metabolism of xenobiotics by cytochrome P450”. These were among the most significantly enriched pathways in both conditions and were predominantly composed of downregulated genes. The consistent predominance of downregulated genes, mainly GST and ADH, within these xenobiotic-metabolism pathways suggests that early exposure to *Esteya* is accompanied by transcriptional downregulation of genes associated with xenobiotic metabolism. This contrasts with the predominantly inducible xenobiotic-detoxification responses reported in *B. xylophilus* under chemical nematicide stress [30,33,35,36] and with the upregulated cytochrome P450-associated response reported following exposure to host-derived monoterpenes [32]. However, downregulation of xenobiotic-metabolism genes is not unique to fungal exposure: a recent transcriptomic study of *B. xylophilus* exposed for 48 h to a lethal concentration of the nematicidal 1,2,4-oxadiazole compound B10 (approximately 1.7-fold its median lethal concentration, LC_50_) also found a predominantly downregulated transcriptional profile, with cytochrome P450- and xenobiotic-metabolism pathways among the enriched categories [37]. Thus, downregulation of xenobiotic-metabolism genes is not diagnostic of fungal challenge. The pattern observed here may nevertheless arise under a different physiological context, since a similar pattern measured after prolonged exposure at a lethal concentration is difficult to separate from a general decline in transcriptional activity, whereas the profile reported here was obtained within 1 h of contact with a live *Esteya* antagonist, under non-lethal exposure conditions.

The higher expression of DNA replication and repair genes under EF represents a further axis of divergence between the two interactions. This suggests that, despite a shared xenobiotic-metabolism response, the two fungi elicited distinct transcriptional programs extending beyond detoxification-associated processes. The functional significance of these differences during the earliest stages of fungal interaction remains unclear and warrants further investigation. One potential explanation for these divergent transcriptional profiles involves species-specific evolutionary histories, though this remains speculative given our single-isolate experimental design. Because fungal virulence was not quantified in parallel, these transcriptional differences cannot be interpreted as evidence of greater host resistance or greater fungal pathogenicity. The broader transcriptional response to EF is consistent with a history of tighter host– pathogen interaction, potentially arising from sympatric co-evolution in North America, the native range of both the PWN and EF [18,38]. While *B. xylophilus* carries an expanded genomic repertoire of detoxification gene families, including cytochrome P450s and UGTs [39], these pathways were transcriptionally downregulated rather than induced during early *Esteya* exposure, suggesting that the early response does not involve induction of this detoxification capacity. By contrast, the more limited response to EV, represented here by a Czech isolate [40], may reflect reduced or delayed host responses, resulting in a narrower transcriptional profile largely nested within that elicited by EF. This interpretation requires validation across multiple isolates of each species and geographic populations, as our comparison of single EF and EV isolates cannot distinguish species-level from isolate-specific effects. While EV isolates from different geographic regions have been characterized, EF is represented by a single described isolate, limiting our ability to assess intraspecific variation for this species. That isolate-level variation can be substantial is not merely theoretical: a quantitative proteomic comparison of two *B. xylophilus* isolates differing in virulence, host, and geographic origin revealed extensive intraspecific divergence, with hundreds of secreted and somatic proteins differentially represented between them [41]. The mixed-stage nematode populations used in our experiment may have introduced developmental variation that could influence baseline transcriptional responses. Furthermore, spore adhesion, germination, and penetration were not quantified microscopically at the time of sampling, so the 1 hpi time point is defined operationally rather than by a verified developmental stage. It therefore cannot be excluded that the observed transcriptional differences partly reflect differences in infection kinetics, particularly relevant for EF, whose infection kinetics have not been described, rather than distinct host responses to the two species. However, the absence of significantly enriched GO categories in the direct EV–EF comparison is consistent with the two fungi having reached broadly comparable stages of interaction at 1 hpi. In addition, because BC served as the fungal-exposure control, the transcriptomic contrasts presented here represent responses to *Esteya* relative to the nematode’s routine laboratory fungal feeding condition, rather than relative to a fungus-free state. Consequently, some of the observed transcriptional changes, particularly those affecting metabolic pathways, may reflect both responses associated with nematophagous fungal exposure and broader physiological adjustments associated with the transition from the standard feeding substrate, and the direction of these changes relative to a fungus-free baseline cannot be established. Additionally, our 1-hour sampling window, while appropriate for capturing early host responses, does not predict longer-term infection outcomes or biocontrol efficacy under field conditions. Future studies incorporating additional EV isolates, defined nematode life stages, and extended time courses will be essential for translating these mechanistic insights into practical biocontrol applications.

From a biocontrol perspective, these findings suggest that early transcriptional signatures may eventually prove useful for predicting interaction outcomes, though the relationship between broader early transcriptional responses and long-term pathogen success remains unclear. Moving forward, understanding inducible cuticle remodeling and other divergent early transcriptional responses may generate hypotheses for selecting or engineering improved *Esteya* strains, although linking these early transcriptional responses to biocontrol performance will require paired virulence assessments. Together, these findings highlight that early interaction dynamics are likely to influence downstream biocontrol outcomes. Disentangling structural versus chemical modes of engagement may therefore be key to selecting and engineering more effective nematophagous fungi.

## Methods

### Biological material and culturing conditions

*Esteya vermicola* CCF3117 [40] and *E. floridanum* V14639 [18] were obtained from culture collections in the Czech Republic and USA, respectively. The reference isolate of *B. xylophilus* Bx013.003 from the Nematology laboratory at the Instituto Nacional de Investigação Agrária e Veterinária (INIAV) was used in this study and reared on a non-sporulating strain of *Botrytis cinerea*. All fungi were grown on potato dextrose agar (PDA, BD Difco™) and incubated at 25 °C for 7 d.

### Sample preparation for transcriptomic analysis

The PWN inoculum was produced as previously described [25] and nematodes extracted using the tray method [42]. PDA plates (90 mm) containing *Esteya* were incubated at 25 °C and further selected based on visual assessment of comparable size and biomass, with inoculation dates staggered to synchronize fungal developmental stage at the time of nematode exposure, as previously observed [25]. This approach ensured that nematodes were exposed to broadly comparable fungal conditions. For each experimental condition, *Esteya* colonies were inoculated with approximately 12,500 mixed-stage nematodes (estimated by extrapolation from nematode density counts in 1 mL using a stereomicroscope at 40× magnification). To capture the early molecular response phase, nematodes were recovered from the fungal mycelium at exactly 1 hpi. Recovery was performed by washing the plates with sterile dH_2_O, gently shaking them by hand and collecting the nematodes in 15 mL Falcon tubes [43]. The recovered suspensions were centrifuged at 3,500 rpm for 5 min and the nematode pellets were transferred to 1.5 mL Eppendorf tubes and immediately frozen at -80 °C. Four biological replicates were used per condition, with the PWN cultured on *B. cinerea* (BC), the standard laboratory rearing fungus on which the nematode feeds and reproduces, serving as the fungal-exposure control. Comparisons with BC therefore represent responses to nematophagous fungi relative to a fungus routinely consumed by the PWN, rather than relative to a fungus-free condition.

### RNA extraction, library preparation and sequencing

Total RNA was extracted from nematodes exposed to each condition using the RNeasy^®^ Mini Kit (QIAGEN, Germany), according to manufacturer’s instructions. DNA digestion was performed with the TURBO DNA-free™ Kit (Thermo Fisher Scientific, USA), to remove residual DNA. RNA integrity was checked by 1% agarose gel electrophoresis. RNA concentrations were measured with both the Invitrogen Qubit 4 Fluorometer (Thermo Fisher Scientific, USA) and NanoDrop™ 2000 Spectrophotometer (Thermo Scientific, USA). Samples were sent to Novogene (Cambridge, UK), where RNA integrity was further assessed on an Agilent 2100 Bioanalyzer (Agilent Technologies, USA); all libraries passed quality control, with RNA integrity numbers of 7.3–9.9. Messenger RNA was isolated from total RNA with poly-T oligo-attached magnetic beads and fragmented, and first-strand cDNA was synthesized with random hexamer primers, followed by second-strand cDNA synthesis. Non-stranded libraries were completed by end repair, A-tailing, adapter ligation, size selection, amplification and purification, quantified by Qubit and real-time PCR, and checked for size distribution on a Bioanalyzer. Libraries were then pooled according to effective concentration and target data output and sequenced on an Illumina NovaSeq X Plus platform in paired-end 150 bp mode, generating a mean of 124.0 million clean reads per sample (range 78.7–180.7 million).

### Raw read mapping and differential gene expression

Raw read processing and bioinformatic analyses were performed by Novogene as part of the RNA-sequencing service. Raw reads were quality-filtered by removing adapter-contaminated reads, reads containing >10% ambiguous bases (N), and low-quality reads in which more than 50% of bases had a Phred quality score ≤5, yielding clean reads for downstream analyses. Clean reads were aligned to the *B. xylophilus* reference genome (PRJEA64437; [44]), available through the National Center for Biotechnology Information (NCBI; https://www.ncbi.nlm.nih.gov/home/genomes/) and WormBase ParaSite release WBPS18 (https://parasite.wormbase.org/), using HISAT2 v2.2.1. Total mapping rates ranged from 86.3% to 91.1%, with 83.6–89.1% of reads mapping uniquely (Supplementary Table S1). Gene expression was quantified with featureCounts v2.0.6 as both raw read counts and fragments per kilobase of transcript per million mapped reads (FPKM). Differential gene expression analysis was performed using raw read counts, which were normalized within the DESeq2 v1.42.0 statistical framework implemented in the Novogene analysis pipeline. Log_2_ fold changes and Benjamini–Hochberg-adjusted *P*-values were used to identify differentially expressed genes. Differentially expressed genes (DEGs) were defined as those with |log_2_FC| ≥ 1 and an adjusted *P*-value (*P_adj_*) ≤ 0.05. Differential expression was assessed separately for each fungal treatment against the BC control (EF vs BC and EV vs BC). Because the two *Esteya* species were never tested together, EF and EV responses were compared in two ways: by contrasting the independent EF vs BC and EV vs BC results, and by a direct DESeq2 contrast between the separately exposed EV and EF samples (EV vs EF). Both approaches describe differences in how the PWN responded to each fungus, rather than the outcome of a co-exposure experiment. Functional enrichment analysis of DEGs was performed with clusterProfiler v4.8.1 using Gene Ontology (GO) and Kyoto Encyclopedia of Genes and Genomes (KEGG) databases to identify significantly overrepresented biological functions and pathways [45–47]. GO annotations were classified into Biological Process, Molecular Function, and Cellular Component categories, and GO and KEGG terms with *P_adj_* < 0.05 were considered significantly enriched.

### Validation of differential gene expression by RT-qPCR

Seven DEGs were selected for RT-qPCR validation, representing genes shared between multiple treatments as well as genes specific to individual contrasts. Gene-specific primers were designed using Primer-BLAST (NCBI) based on *B. xylophilus* reference gene models (Table 1). Total RNA was reverse-transcribed into cDNA using the SuperScript™ IV First-Strand Synthesis System (Thermo Fisher Scientific, USA) according to the manufacturer’s instructions. Before RT-qPCR, each primer pair was tested by conventional PCR on *B. xylophilus* genomic DNA and cDNA to confirm amplification of the target transcripts, and amplicons were checked by agarose gel electrophoresis. Amplification was performed using a SYBR Green assay with the NZYSupreme qPCR Green Master Mix (NZYtech, Lisbon, Portugal) under the following thermal cycling conditions: initial denaturation at 95 °C for 3 min, followed by 40 cycles of 95 °C for 20 s and 60 °C for 30 s (55 °C for BXY_0475400), and a melting curve analysis. A single melting peak for each amplicon confirmed primer specificity. No-template controls were included in each run. Transcript abundance was normalized to the reference gene actin (*Bxy-act-1*) [48], and relative expression levels were calculated using the 2^−ΔΔCT^ method [49]. For RT-qPCR validation, each sample was measured in two or three technical replicates.

**Table 1.**
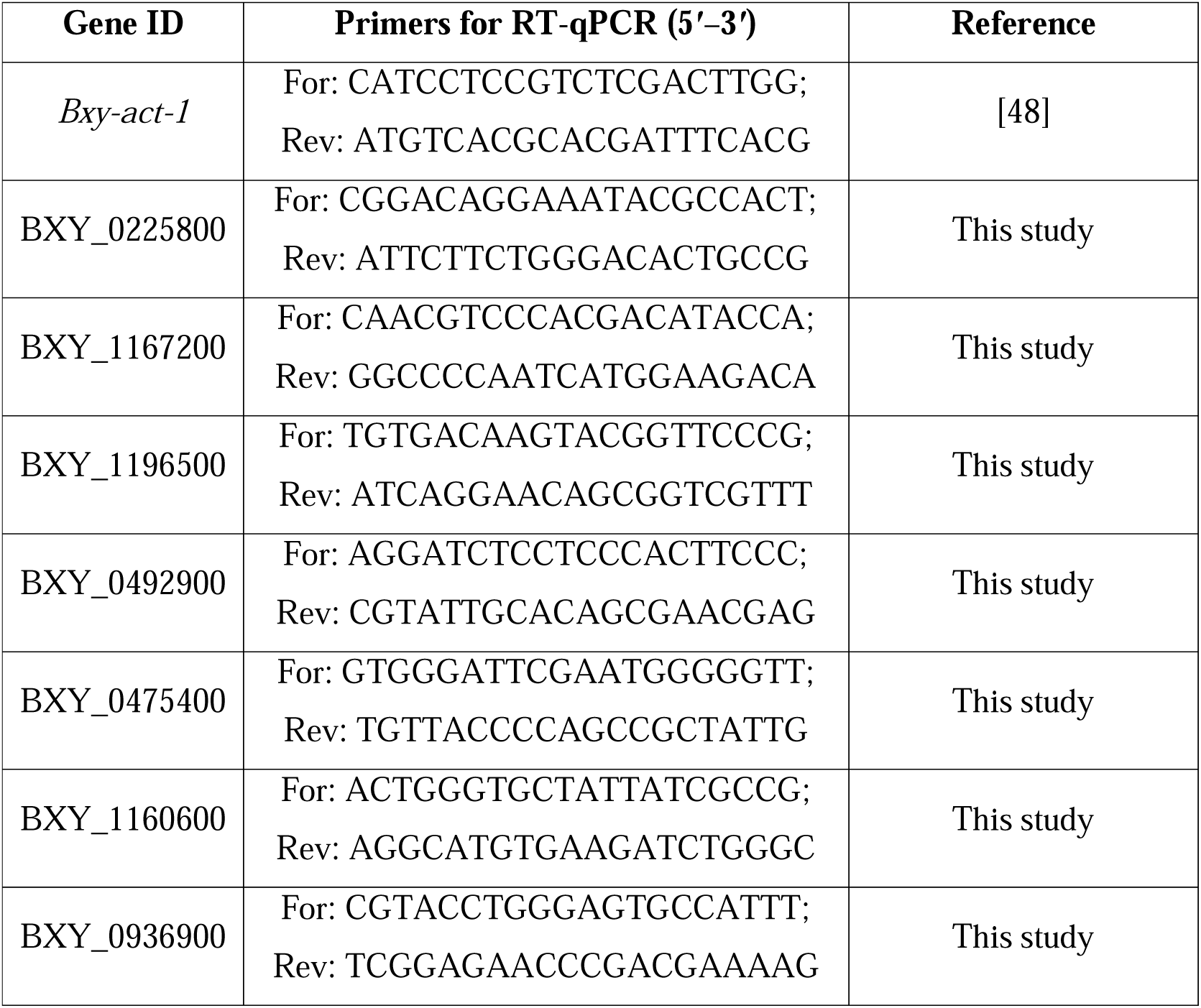
Genes selected for RT-qPCR validation and primer sequences used for amplification. The reference gene *Bxy-act-1* was used for normalization of transcript abundance.

### Data visualization

Visualization of results, including volcano plots, GO and KEGG enrichment dot plots, and Venn diagrams, was provided by Novogene (Cambridge, UK) as part of their comprehensive transcriptomic analysis pipeline. The petal diagram was recreated in Adobe Photoshop CC based on the original Venn diagram output from Novogene, with colors adjusted to visually match the graphical abstract design.

### Use of large language models

The authors used Claude Opus 5 (Anthropic) to improve language and for proofreading. All content was subsequently reviewed and edited by the authors, who take full responsibility for the final version.

## Data availability

The RNA-seq datasets generated and analyzed during this study are available in the National Center for Biotechnology Information (NCBI) database under BioProject accession PRJNA1519337 (https://www.ncbi.nlm.nih.gov/bioproject/PRJNA1519337). The raw sequencing reads are available through the Sequence Read Archive (SRA) under accession numbers SRX35015406–SRX35015417.

## Acknowledgements

This work was conducted within the framework of an international collaboration established during DP’s doctoral research between the Instituto Nacional de Investigação Agrária e Veterinária (INIAV), the Mediterranean Institute for Agriculture, Environment and Development (MED), and Charles University. We gratefully acknowledge Alena Kubátová (Charles University) for facilitating this collaboration and DP’s research stay in the Czech Republic. The authors express their gratitude to the members of the Nematology laboratories at INIAV and MED–University of Évora for their assistance and constructive discussions. We also thank the members of the Biochemistry laboratory at INIAV for their support.

## Author contributions

**David Pires**: Conceptualization, Data curation, Formal analysis, Investigation, Methodology, Validation, Visualization, Writing – Original Draft, Writing – Reviewing and Editing. **Ana Rita Varela**: Formal analysis, Validation, Writing – Reviewing and Editing. **Miroslav Kola**ř**ík**: Writing – Reviewing and Editing. **Maria L. Inácio**: Funding acquisition, Writing – Reviewing and Editing. **Manuel Mota**: Writing – Reviewing and Editing. **Cláudia S. L. Vicente**: Conceptualization, Data curation, Formal analysis, Funding acquisition, Methodology, Supervision, Validation, Writing – Reviewing and Editing.

## References

1. Zhou, J. et al. Vulnerability of global pine forestry’s carbon sink to an invasive pathogen–vector system. Glob. Chang. Biol. 30, e17614 (2024).

2. Leroy, B. M. L. Global insights on insecticide use in forest systems: current use, impacts and perspectives in a changing world. Curr. For. Rep. 11, 6 (2024).

3. Sun, H. et al. How to eradicate an invasive forest pest without clear cutting. J. Appl. Ecol. 63, e70318 (2026).

4. EFSA Panel on Plant Health et al. Systematic review of modelling approaches assessing eradication and containment measures for the pinewood nematode (PWN) Bursaphelenchus xylophilus. EFSA J. 24, (2026).

5. Hoffmann, V. et al. A one health approach to plant health. CABI Agric. Biosci. 3, 62 (2022).

6. Schaffner, U. et al. Biological control for One Health. Sci. Total Environ. 951, 175800 (2024).

7. Borgianni, L., et al. The power of microbial life for the transformation towards a sustainable planet: key messages from the 2024 IUMS Congress in Florence, the city of the Renaissance. microLife 6, (2025).

8. Back, M. A., Bonifácio, L., Inácio, M. L., Mota, M. & Boa, E. Pine wilt disease: a global threat to forestry. Plant Pathol. 73, 1026–1041 (2024).

9. Akira, S., Uematsu, S. & Takeuchi, O. Pathogen recognition and innate immunity. Cell 124, 783–801 (2006).

10. Pujol, N. et al. Anti-fungal innate immunity in *C. elegans* is enhanced by evolutionary diversification of antimicrobial peptides. PLoS Pathog. 4, e1000105 (2008).

11. Gravato-Nobre, M. J. & Hodgkin, J. *Caenorhabditis elegans* as a model for innate immunity to pathogens. Cell. Microbiol. 7, 741–751 (2005).

12. Chang, H.-W. et al. Cuticular collagens mediate cross-kingdom predator–prey interactions between trapping fungi and nematodes. PLoS Biol. 23, e3003178 (2025).

13. Pires, D., Vicente, C. S. L., Inácio, M. L. & Mota, M. The potential of *Esteya* spp. for the biocontrol of the pinewood nematode, *Bursaphelenchus xylophilus*. Microorganisms 10, 168 (2022).

14. Pires, D., Vicente, C. S. L., Mota, M. & Inácio, M. L. *Esteya* spp. reduce *Bursaphelenchus xylophilus* reproduction in *Pinus pinaster* seedlings without consistent improvements in disease outcomes. Pest Manag. Sci. 10.1002/ps.71200 (2026).

15. Liou, J. Y., Shih, J. Y. & Tzean, S. S. *Esteya*, a new nematophagous genus from Taiwan, attacking the pinewood nematode (*Bursaphelenchus xylophilus*). Mycol. Res. 103, 242–248 (1999).

16. Wang, C. Y. et al. High infectivity of an endoparasitic fungus strain, *Esteya vermicola*, against nematodes. J. Microbiol. 46, 380–389 (2008).

17. Wang, C. Y. et al. High infection activities of two *Esteya vermicola* isolates against pinewood nematode. Afr. J. Microbiol. Res. 3, 581–584 (2009).

18. Li, Y., et al. *Esteya floridanum* sp. nov.: an ophiostomatalean nematophagous fungus and its potential to control the pine wood nematode. Phytopathology 111, 304–311 (2021).

19. Wang, X. et al. Isolation and identification of *Esteya vermicola* and its potential for controlling pinewood nematode. For. Pathol. 52, e12745 (2022).

20. Wang, H. H. et al. In vivo infection of *Bursaphelenchus xylophilus* by the fungus *Esteya vermicola*. Pest Manag. Sci. 76, 2854–2864 (2020).

21. Lin, F., Ye, J., Wang, H., Zhang, A. & Zhao, B. Host deception: predaceous fungus, *Esteya vermicola*, entices pine wood nematode by mimicking the scent of pine tree for nutrient. PLoS One 8, e71676 (2013).

22. Wang, X., Guan, T., Zhang, L. & Li, H. Cloning of a serine protease gene from the nematophagous fungus *Esteya vermicola* and expressed activity of the recombinant enzyme against *Bursaphelenchus xylophilus*. Nematology 17, 1071– 1080 (2015).

23. Wang, R., et al. *Esteya vermicola*, a nematophagous fungus attacking the pine wood nematode, harbors a bacterial endosymbiont affiliated with Gammaproteobacteria. Microbes Environ. 32, 201–209 (2017).

24. Wang, R., Dong, L., Chen, Y., Wang, S. & Qu, L. Third generation genome sequencing reveals that endobacteria in nematophagous fungi *Esteya vermicola* contain multiple genes encoding for nematicidal proteins. Front. Microbiol. 13, 842684 (2022).

25. Pires, D., Vicente, C. S. L., Mota, M. & Inácio, M. L. Polyphasic approach to the selection of *Esteya* isolates for the control of the pinewood nematode, *Bursaphelenchus xylophilus*. Fungal Biol. 128, 2242–2249 (2024).

26. Hammond, R., Dickinson, A., Damm, A. & Eves-van den Akker, S. The five senses: how do plant pathogens know they found their host? Mol. Plant Microbe Interact. 39, 4–9 (2026).

27. Espada, M., Vicente, C. S. L. & Vieira, P. Molecular insights into migratory plant-parasitic nematodes. Annu. Rev. Phytopathol. 63, 403–430 (2025).

28. Hsueh, Y.-P., Mahanti, P., Schroeder, F. C. & Sternberg, P. W. Nematode-trapping fungi eavesdrop on nematode pheromones. Curr. Biol. 23, 83–86 (2013).

29. Kuo, C.-Y. et al. The nematode-trapping fungus *Arthrobotrys oligospora* detects prey pheromones via G protein-coupled receptors. Nat. Microbiol. 9, 1738–1751 (2024).

30. Hao, X. et al. Molecular defense response of *Bursaphelenchus xylophilus* to the nematophagous fungus *Arthrobotrys robusta*. Cells 12, 543 (2023).

31. Xiong, X. et al. BxALDH as a potential target of fluensulfone for controlling *Bursaphelenchus xylophilus* in industrial pine trees. Ind. Crops Prod. 250, 123835 (2026).

32. Duan, W., Zhang, X., Duan, J., Sun, J. & Zhang, B. The pinewood nematode enhances population growth by exploiting host defensive terpenes via lysosomal signaling. Pest Manag. Sci. 10.1002/ps.71074 (2026).

33. Chen, J., Hao, X., Wang, B. & Ma, L. Transcriptomics and coexpression network profiling of the effects of levamisole hydrochloride on *Bursaphelenchus xylophilus*. Pestic. Biochem. Physiol. 181, 105019 (2022).

34. Yu, S. et al. Nematicidal activity of 3-indoleacetonitrile against *Bursaphelenchus xylophilus*: Inhibition of development and reproduction through induction of excessive reactive oxygen species. Pestic. Biochem. Physiol. 213, 106509 (2025).

35. Li, Y. et al. Functional analysis of 3 genes in xenobiotic detoxification pathway of *Bursaphelenchus xylophilus* against matrine. Pestic. Biochem. Physiol. 190, 105334 (2023).

36. Cao, J. et al. Exploring the role of detoxification genes in the resistance of *Bursaphelenchus xylophilus* to different exogenous nematicidal substances using transcriptomic analyses. Pestic. Biochem. Physiol. 194, 105527 (2023).

37. Zhao, R., et al. Discovery of double 1,2,4 oxadiazole derivatives as promising nematicides. Pest Manag. Sci. 10.1002/ps.71151 (2026).

38. Tares, S., Lemontey, J. M., de Guiran, G. & Abad, P. Use of species-specific satellite DNA from *Bursaphelenchus xylophilus* as a diagnostic probe. Phytopathology 84, 294–298 (1994).

39. Xiong, X. et al. Genome-wide characterization of the *Bursaphelenchus xylophilus* UGT gene family and its potential roles in detoxification and host interaction. Front. Plant Sci. 17, 1786843 (2026).

40. Kubátová, A., Novotný, D., Prášil, K. & Mráček, Z. The nematophagous hyphomycete *Esteya vermicola* found in the Czech Republic. Czech Mycol. 52, 227–235 (2000).

41. Cardoso, J. M. S. et al. Virulence biomarkers of *Bursaphelenchus xylophilus*: a proteomic approach. Front. Plant Sci. 12, 822289 (2022).

42. Whitehead, A. G. & Hemming, J. R. A comparison of some quantitative methods of extracting small vermiform nematodes from soil. Ann. Appl. Biol. 55, 25–38 (1965).

43. Lin, H.-C. et al. Key processes required for the different stages of fungal carnivory by a nematode-trapping fungus. PLoS Biol. 21, e3002400 (2023).

44. Kikuchi, T. et al. Genomic insights into the origin of parasitism in the emerging plant pathogen *Bursaphelenchus xylophilus*. PLoS Pathog. 7, e1002219 (2011).

45. Ashburner, M. et al. Gene Ontology: tool for the unification of biology. Nat. Genet. 25, 25–29 (2000).

46. Kanehisa, M. & Goto, S. KEGG: Kyoto Encyclopedia of Genes and Genomes. Nucleic Acids Res. 28, 27–30 (2000).

47. Aleksander, S. A. et al. The Gene Ontology knowledgebase in 2023. Genetics 224, iyad031 (2023).

48. Vicente, C. S. L., Ikuyo, Y., Shinya, R., Mota, M. & Hasegawa, K. Catalases induction in high virulence pinewood nematode *Bursaphelenchus xylophilus* under hydrogen peroxide-induced stress. PLoS One 10, e0123839 (2015).

49. Livak, K. J. & Schmittgen, T. D. Analysis of relative gene expression data using real-time quantitative PCR and the 2^−ΔΔCT^ method. Methods 25, 402–408 (2001).

