## Supplementary Information for "Sister species, shared and divergent responses: early transcriptomic profiling of *Bursaphelenchus xylophilus* exposed to *Esteya* spp"

**Supplementary material**

**Table S1**. Sequencing and mapping statistics for the twelve RNA-seq libraries.

| **Sample** | **Raw**  **reads** | **Clean**  **reads** | **Raw bases**  **(Gb)** | **Clean bases**  **(Gb)** | **Q20**  **(%)** | **Q30**  **(%)** | **GC**  **(%)** | **Total**  **mapping** | **Unique**  **mapping** |
| --- | --- | --- | --- | --- | --- | --- | --- | --- | --- |
| EV_1 | 136,289,122 | 132,739,722 | 20.44 | 19.91 | 99.42 | 97.04 | 48.61 | 115,506,553 (87.02%) | 111,641,418 (84.11%) |
| EV_2 | 101,624,938 | 98,023,258 | 15.24 | 14.70 | 99.39 | 97.03 | 48.21 | 86,595,473 (88.34%) | 84,742,179 (86.45%) |
| EV_3 | 107,519,062 | 104,395,226 | 16.13 | 15.66 | 99.38 | 97.02 | 48.48 | 91,456,891 (87.61%) | 89,119,965 (85.37%) |
| EV_4 | 138,066,154 | 134,037,844 | 20.71 | 20.11 | 99.43 | 97.02 | 48.20 | 116,957,309 (87.26%) | 114,088,114 (85.12%) |
| EF_1 | 149,742,876 | 145,586,406 | 22.46 | 21.84 | 99.41 | 97.03 | 48.57 | 125,607,470 (86.28%) | 121,631,018 (83.55%) |
| EF_2 | 100,911,342 | 96,678,598 | 15.14 | 14.50 | 99.38 | 96.85 | 48.45 | 84,084,092 (86.97%) | 81,229,891 (84.02%) |
| EF_3 | 161,950,264 | 159,244,794 | 24.29 | 23.89 | 99.42 | 96.98 | 48.19 | 143,863,409 (90.34%) | 140,500,721 (88.23%) |
| EF_4 | 185,382,902 | 180,682,206 | 27.81 | 27.10 | 99.43 | 97.04 | 48.01 | 160,737,770 (88.96%) | 157,616,322 (87.23%) |
| BC_1 | 132,949,606 | 129,044,010 | 19.94 | 19.36 | 99.39 | 96.89 | 48.88 | 113,474,397 (87.93%) | 109,724,778 (85.03%) |
| BC_2 | 99,951,082 | 96,824,446 | 14.99 | 14.52 | 99.40 | 97.00 | 47.94 | 88,170,480 (91.06%) | 86,277,649 (89.11%) |
| BC_3 | 81,973,616 | 78,736,926 | 12.30 | 11.81 | 99.39 | 97.02 | 48.57 | 70,497,660 (89.54%) | 69,094,857 (87.75%) |
| BC_4 | 138,608,768 | 132,484,264 | 20.79 | 19.87 | 99.40 | 96.90 | 48.69 | 117,618,068 (88.78%) | 114,677,776 (86.56%) |

EV, *Esteya vermicola*; EF, *E. floridanum*; BC, *Botrytis cinerea*. Mapping values are given as read counts with the corresponding percentage of clean reads.

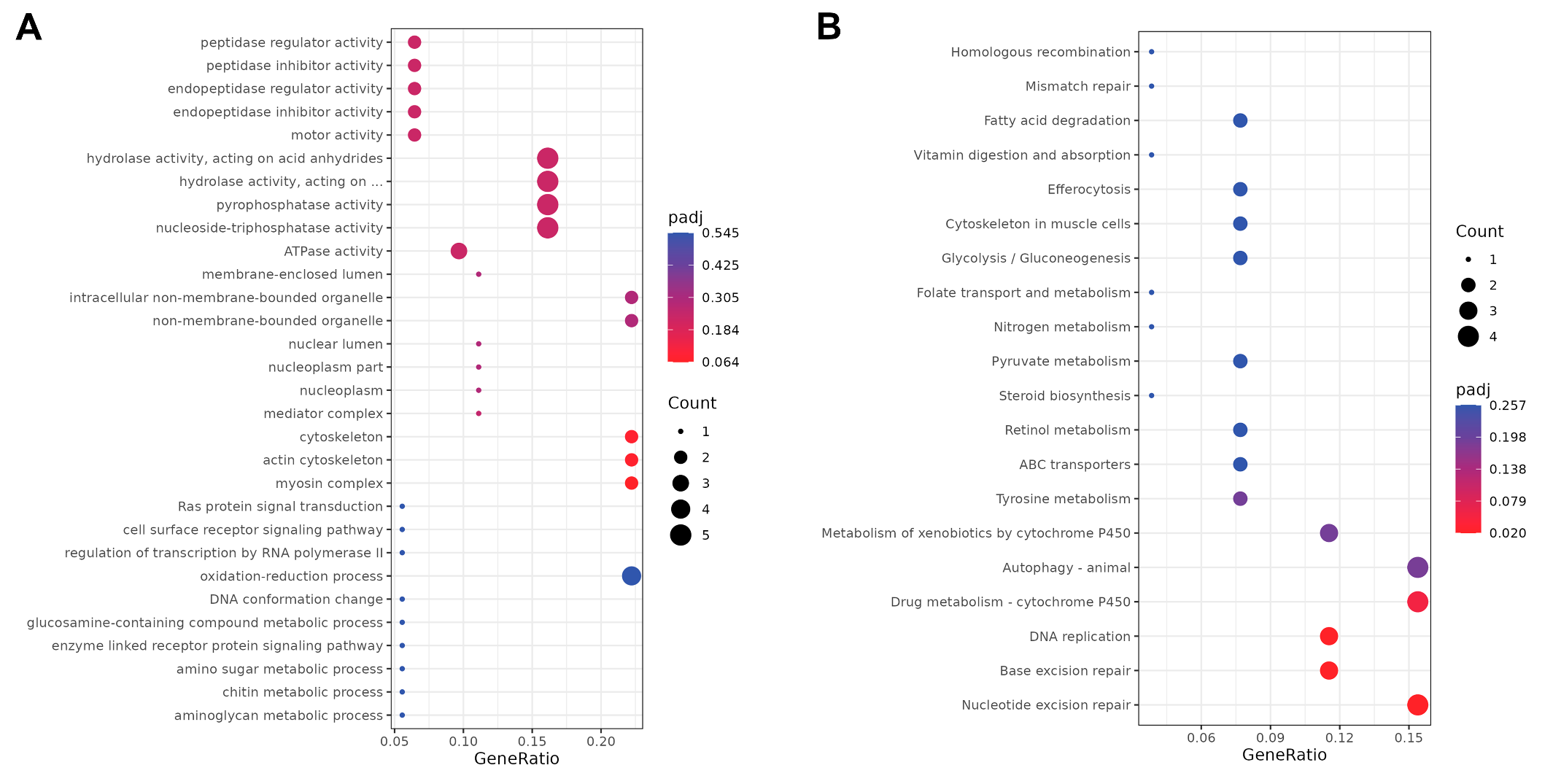

**Figure S1**. Functional enrichment of the direct comparison between Bursaphelenchus xylophilus exposed to E. vermicola and E. floridanum (EV vs EF). (A) Top-ranked Gene Ontology (GO) terms, none of which were significantly enriched. (B) Top-ranked KEGG pathways; nucleotide excision repair, base excision repair and DNA replication were significantly enriched (P_adj_ = 0.020). Dot size represents the number of DEGs annotated to each term or pathway, and color represents the adjusted P-value; color scales differ between panels.

**Table S2.** RT-qPCR validation of selected differentially expressed genes.

| **Gene ID** | **Annotation** | **Contrast** | **RNA-seq log_2_FC** | **RNA-seq *P*_adj_** | **RT-qPCR log_2_FC**  **(mean ± SD)** | **Concordance** |
| --- | --- | --- | --- | --- | --- | --- |
| BXY_0225800 | Glucuronosyltransferase  (*sqv-8* homologue) | EV vs BC | 3.28 | 2.8 × 10^−3^ | 2.05 ± 1.21 | Yes |
|  |  | EF vs BC | 3.81 | 3.9 × 10^−9^ | −0.41 ± 1.15 | No |
| BXY_1167200 | Flavin-containing  monooxygenase | EV vs BC | −2.68 | 1.8 × 10^−29^ | −3.02 ± 0.77 | Yes |
|  |  | EV vs EF | −2.64 | 5.0 × 10^−15^ | −4.99 ± 0.65^†^ | Yes |
| BXY_0936900 | Hypothetical protein | EF vs BC | 5.14 | 1.8 × 10^−2^ | 3.91 ± 1.11 | Yes |
| BXY_1160600 | Hypothetical protein | EF vs BC | −6.96 | 1.4 × 10^−7^ | 2.68 ± 1.77 | No |
| BXY_1196500 | Hypothetical protein | EV vs EF | 1.74 | 6.0 × 10^−3^ | −2.50 ± 0.33^†^ | No |
| BXY_0492900 | Alcohol dehydrogenase  (*sodh-2* homologue) | EV vs EF | 1.23 | 4.0 × 10^−3^ | 0.40 ± 1.14^†^ | Yes |
| BXY_0475400 | Hypothetical protein | EV vs BC | −4.87 | 2.9 × 10^−13^ | −0.39 ± 2.13^‡^ | Yes |
|  |  | EF vs BC | −4.18 | 1.2 × 10^−13^ | −2.04 ± 1.85^‡^ | Yes |

RT-qPCR log_2_FC = −ΔΔCt (Livak & Schmittgen, 2001), using *Bxy-act-1* as reference gene and *B. xylophilus* cultured on *B. cinerea* (BC) as calibrator. Values are means ± SD of four biological replicates per group (except BXY_1196500, EV *n* = 3; BXY_0492900, EF *n* = 3), each measured in two or three technical replicates. RNA-seq values are DESeq2 log_2_ fold changes and Benjamini–Hochberg-adjusted *P*-values. Concordance indicates the same direction of change in both methods. ^†^EV vs EF calculated as log_2_FC(EV/BC) − log_2_FC(EF/BC); the error is the standard error of the difference. ^‡^Low-abundance transcript (Ct > 30). Pearson’s correlation between RNA-seq and RT-qPCR log_2_FC across the ten quantified contrasts: *r* = 0.30 (*P* = 0.41). EV, *Esteya vermicola*; EF, *E. floridanum*.

.
